# Impact of dopaminergic modulation on the state transitions of striatal medium spiny neuron sub-types – a computational study

**DOI:** 10.1101/2024.12.22.629989

**Authors:** Nitin Anisetty, Rohit Manchanda

## Abstract

Medium spiny neurons (MSN) of the striatum are known for their bistable membrane potential leading to two states: a hyperpolarized down-state and a depolarized up-state. Glutamatergic inputs from the hippocampus play a key role in switching the cell to the up-state. This gating is known to play a key role in regulating when other synaptic inputs, such as from the cortex, should generate action potentials and when they should be considered as noise. Any deviations from this pattern of state transitions indicates abnormal gating that has implications in conditions such as schizophrenia. Although dopamine is reported to modulate ion channels of MSN sub-types - dMSN and iMSN - its influence on the state transition times and up-state dwell times are not yet examined. We address this lacuna using biophysically constrained spiny models of dMSN and iMSN with explicit dopamine receptors. Our findings indicate a significant increase in up-state dwell time for dMSN and a significant decrease for iMSN when the % activation of DA receptors was increased. Additionally, a strong correlation between state transition times and spiking frequencies of MSN sub-types was observed.

## Introduction

The basal ganglia (BG) are a group of nuclei in the brain that are primarily associated with motor control and value-based action selection. The striatum is the key input region of BG that predominantly constitutes cells called Medium spiny neurons (MSN). The MSN are crucial in driving two major pathways of the BG owing to their high computational capacity to process multiple excitatory inputs from regions such as the hippocampus, amygdala, prefrontal cortex, and thalamus, as well as inhibitory inputs from MSN lateral connections and GABAergic interneurons (Li et al., 2018). MSN can be classified into dMSN and iMSN. The dMSN typically express D1 dopamine receptor, whereas the iMSN express D2 dopamine receptor, thereby enabling dopamine (DA) to influence dMSN and iMSN differentially.

The output nuclei of the Basal Ganglia (BG) i.e. SNr (Substantia Nigra pars reticulata) and GPi (Globus pallidus internus) receive axonal projections from the dMSN directly whereas they are influenced indirectly by the iMSN via intermediate regions, namely, GPe (Globus Pallidus externus) and STN (Subthalamic nucleus). These pathways from the striatal neurons to the output nuclei of BG are therefore referred to as the direct pathway and indirect pathway, respectively. The segregation of dMSN and iMSN with respect to direct and indirect pathway typically holds true for dorsal striatum as described and also via analogous structures from the nucleus accumbens core region of the ventral striatum (Gangarossa et al., 2013). Both pathways together contribute in the selection or avoidance of actions. Therefore, the physiological properties of these MSN sub-types are of prime importance to study.

A characteristic feature of MSN is the bi-stability of membrane potential described by several research groups (Wilson and Kawaguchi, 1996; Gruber et al., 2003). Glutamatergic inputs primarily from the hippocampus are said to facilitate the transition of MSN from a hyperpolarized down-state to a depolarized up-state (O’Donnell and Grace, 1995). Hippocampus is known to encode episodic memories (Kolibius et al., 2023), therefore, it can be speculated that the switch to up-state occurs in context of subject recollecting experiences related to the action in question. Furthermore, glutamatergic inputs from other regions such as the pre-frontal cortex do not trigger switch to up-state and action potentials are not triggered in the down-state. Therefore, cortical inputs have greater influence only when contextual information arrives via limbic inputs. The state transitions are consequently crucial and act as filters when the synaptic inputs are relevant as opposed to when they are to be considered as noise. Abnormal gating has implications in the pathophysiology of schizophrenia (Grace, 2000). The up-state duration could vary from 100-1000 ms, although, the average duration was reported to be 284 ms as was simulated in our model by controlling the synaptic input drive (Blackwell et al., 2003).

Dopamine (DA) has been widely reported to signal reward prediction error (RPE) that is encoded in cortico-striatal synapses of MSN (Gurney et al., 2015). However, there is growing evidence for its contribution in other functions (Lau et al., 2017). Apart from regulating long-term synaptic plasticity at cortico-striatal synapses, dopamine is known to transiently modulate ion channels (Nicola et al., 2000). Using biophysically constrained models of dMSN and iMSN with explicit D1R and D2R respectively, we investigate how DA affects the gating of states in dMSN and iMSN. The output measures used to quantify this are the state transition times and the up-state dwell time. Furthermore, we analyse how the state transition times are correlated with the spiking frequency of dMSN and iMSN under different % activation of DA receptors (D1R and D2R).

The models are first-of-its-kind with the dendritic spines and localized dendritic segments integrated with mechanisms for spatiotemporal DA induced ion channel modulation. To validate the models, they were subjected to current injection inputs which, as anticipated from physiological data, indicated a greater intrinsic excitability of the iMSN over dMSN. The results indicated that the transition to upstate was quicker and transition to downstate was slower for dMSN with increasing % DA receptor (D1R) activation. For the iMSN, the direction of change was opposite with increasing % DA receptor (D2R) activation. As anticipated, this indicated a higher up-state dwell time for dMSN and lower up-state dwell time for iMSN with increasing % DA receptor activation. This is crucial as it implies a greater window for spike generation when the up-state dwell time is longer. Although there is a possibility that a greater spiking window does not translate to greater spiking, the results did find that there was a strong correlation between spiking frequency and transition times.

## Methods

### Description of the models

For this study, biophysically constrained models of dMSN and iMSN were considered with reference to the available data. Both dMSN and iMSN models were derived primarily from a computational MSN model proposed by (Wolf et al., 2005), but with the inclusion of dendritic spines modified based on (Mattioni and Le Novere, 2013). Furthermore, the two sub-types – dMSN and iMSN – were explicitly linked to D1 and D2 dopamine receptors, respectively (Le Moine and Bloch, 1995; Gerfen and Surmeier, 2011). In the models, the dopamine receptors are located on each dendritic spine neck as well as the middle and tertiary dendrites which facilitate spatiotemporally varied transient ion channel and synaptic modulation when triggered by dopamine, as quantified in (Moyer et al., 2007). The mechanism for intracellular calcium stores was adapted from (Nakano et al., 2013) with some key unit corrections. All implementations were done using NEURON simulation environment with Python, HOC and NMODL programming languages. The models are currently on ModelDB with private access (https://modeldb.science/2020337). An access code can be provided upon request.

#### Biophysical properties: dMSN and iMSN models

The biophysical differences between dMSN and iMSN are reported to be minimal as they are sub-types of MSN (Gertler et al., 2008; Al-Muhtasib et al., 2018). Only Kir channel conductance differed between dMSN and iMSN. Several MSN computational models were compared and the conductance (permeability in case of calcium) values that were most used across several other published models were finalized (refer Table 1).

**Table 1:**
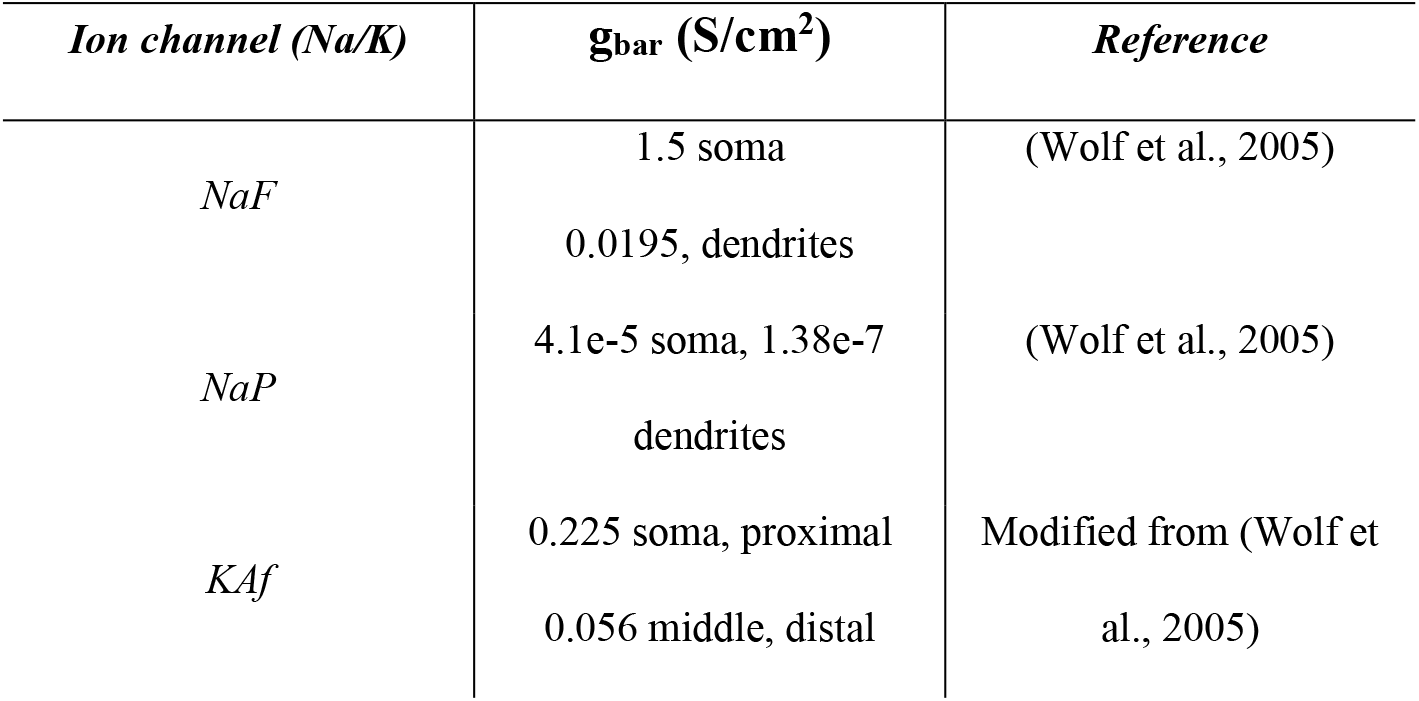
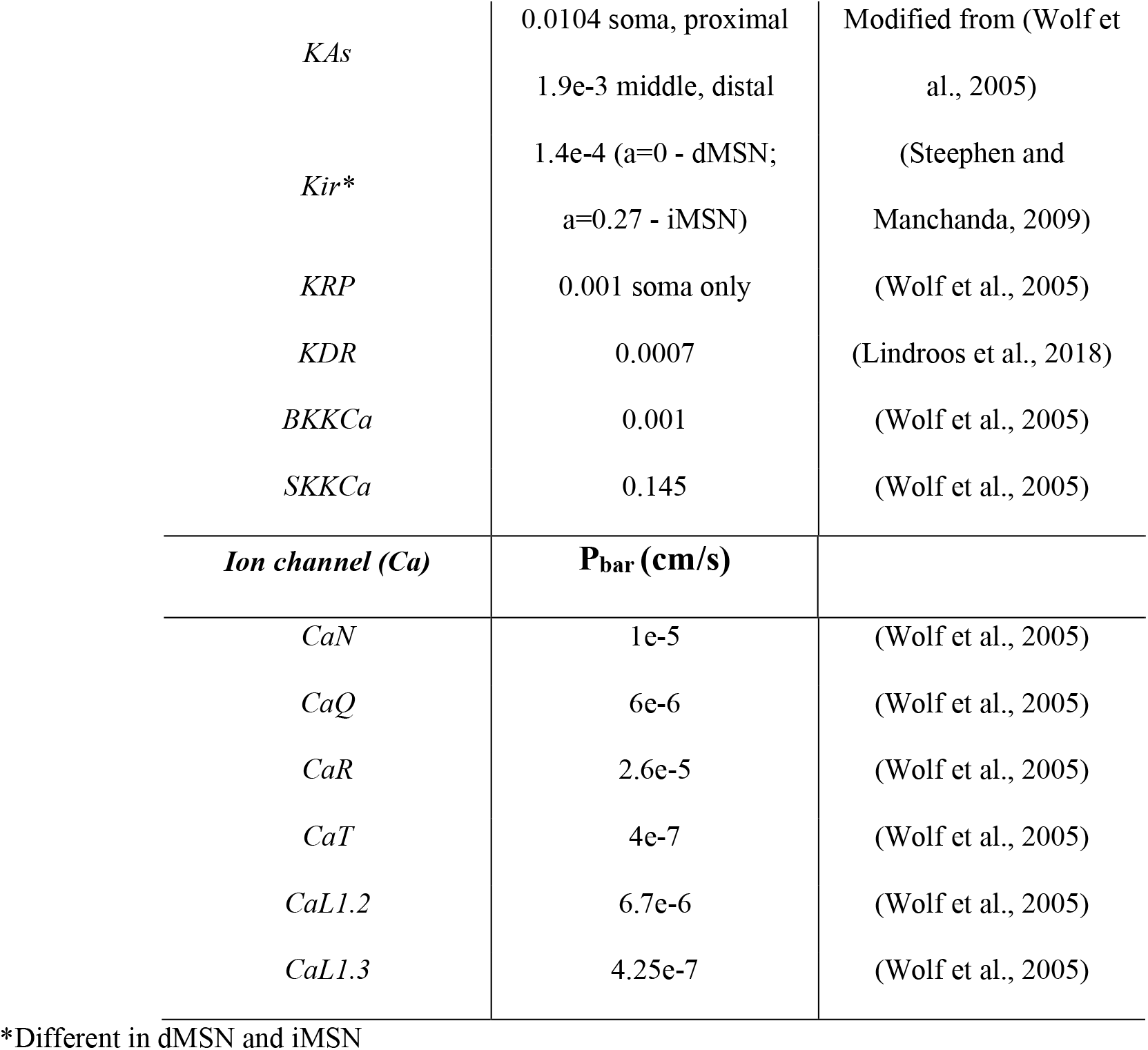
Parameters used for active properties in the model.

| *Ion channel (Na/K)* | *$g_{\text{bar}}$ (S/cm<sup>2</sup>)* | *Reference* |
| --- | --- | --- |
| *NaF* | 1.5 soma | (Wolf et al., 2005) |
|  | 0.0195, dendrites |  |
| *NaP* | 4.1e-5 soma, 1.38e-7 | (Wolf et al., 2005) |
|  | dendrites |  |
| *KAf* | 0.225 soma, proximal | Modified from (Wolf et al., 2005) |
|  | 0.056 middle, distal |  |
| *KAs* | 0.0104 soma, proximal | Modified from (Wolf et |
|  | 1.9e-3 middle, distal | al., 2005) |
| *Kir\** | 1.4e-4 (a=0 - dMSN;<br>a=0.27 - iMSN) | (Steephen and<br>Manchanda, 2009) |
| *KRP* | 0.001 soma only | (Wolf et al., 2005) |
| *KDR* | 0.0007 | (Lindroos et al., 2018) |
| *BKKCa* | 0.001 | (Wolf et al., 2005) |
| *SKKCa* | 0.145 | (Wolf et al., 2005) |
| ***Ion channel (Ca)*** | **$P_{bar}$ (cm/s)** |  |
| *CaN* | 1e-5 | (Wolf et al., 2005) |
| *CaQ* | 6e-6 | (Wolf et al., 2005) |
| *CaR* | 2.6e-5 | (Wolf et al., 2005) |
| *CaT* | 4e-7 | (Wolf et al., 2005) |
| *CaL1.2* | 6.7e-6 | (Wolf et al., 2005) |
| *CaL1.3* | 4.25e-7 | (Wolf et al., 2005) |

\*Different in dMSN and iMSN

Apart from the difference in dopamine receptors expressed, further modifications are indicated as follows:

- AMPA receptors undergo desensitization in the model upon repeated exposure to inputs (Hansen et al., 2007).
- Sign correction of the variable ‘alp’ of BKKCa channel mechanism used in (Wolf et al., 2005) model. The correction is consistent with mechanism used by (Steephen and Manchanda, 2009).
- It has been reported that neuropeptide Enkephalin (ENK) positive neurons have partially inactivating Kir channels while neuropeptide Substance P (SP) positive neurons have non-inactivating Kir channels (Steephen and Manchanda, 2009). Several reports indicate that dMSN contains SP while iMSN contains ENK (Humphries and Prescott, 2010). Therefore, in our model we have used non-inactivating Kir channels for dMSN and partially inactivating Kir channels for iMSN.
- The variable ‘ainh’ in ‘ER’ mechanism (for Ca^+2^ intracellular store) was corrected as incorrect units led to incorrect value in (Nakano et al., 2013). It was 2e-4 mM/s but the corrected value and units is 0.2 1/mM-ms found after backtracking to original source of value: (Li and Rinzel, 1994).

#### Morphological properties: dMSN and iMSN models

The dendritic morphology of these neurons is known to follow a bifurcating pattern, i.e the distal dendrites bifurcate from middle dendrites which in turn bifurcate from proximal dendrites. This was well represented by the stylized morphology first adapted by (Wolf et al., 2005) and then in several models thereafter. Furthermore, iMSN are reported to be inherently more excitable than dMSN, putatively due to smaller total dendritic length. The median total dendritic length of dMSN and iMSN was found to be 3283 μm and 2532 μm, respectively (Gertler et al., 2008). This was attributed to a difference in the number of dendritic branches. Consistent with these findings, we have reduced the branching of iMSN (3 primary, 6 middle and 12 distal dendrites) in comparison to dMSN (4 primary, 8 middle and 16 distal dendrites). This results in the total dendritic length of dMSN coming to 3280 μm and of iMSN coming to 2460 μm, respectively, closely resembling the lengths reported from experimental studies. In accordance with the lower branching of iMSN, the number of glutamatergic and GABAergic inputs were reduced while maintaining their ratio (∼1:1) constant across the two cell types.

Each individual dendritic spine used in our model was considered as a 2-compartment structure consisting of the spine head and the spine neck. This was adapted from Mattioni et al., 2013 but was modified by merging the spine PSD and head section in order to facilitate interaction of ions like calcium via a shared calcium pool between the PSD zone and head region as also implemented in (Nakano et al., 2013). Dendritic spines were incorporated on the middle and distal dendrites. There were 13 spines on each middle dendrite and 307 spines on each distal dendrite. The spines on the distal dendrites were located principally at the proximal end i.e. closer to its connection with the middle dendrite as reported (Wilson et al., 1983; Rane and Manchanda, 2017). Therefore, for each pseudo-randomized generation of the model, we use a gaussian distribution (μ = 85 μm, SD = 25 μm) such that 95% of spines lie between 35 to 135 μm on a 190 μm length distal dendrite. The middle dendrites are small in size and are modelled as a single compartment so the dendritic spines here are spatially similar across trials.

As the focus of the study is the role of DA on the differential excitabilities of dMSN and iMSN via its action on ion channels, apart from the above-mentioned differences, we constrained the effect of morphological differences on the differential excitability. In this manner, the potential variability of excitability from trial-to-trial due to morphological randomness is regulated.

### Modelling dopamine interaction with dMSN and iMSN

Transient dopamine modulation occurs at sub-second timescale, therefore, this hypothesis is best tested in the computational domain using spatiotemporally varied dopaminergic inputs projecting to biophysically constrained computational models of both MSN sub-types (dMSN and iMSN). Intrinsic ion channels in our MSN models are modulated temporally by a variable called dopamine modulation factor (μ) described by (Gruber et al., 2003). Upon activation of a DA receptor (D1R or D2R), μ rises for 60 ms with a time constant (τ_inc_) of 30 ms indicating DA bound to receptor and thereafter decays with a time constant (τ_dec_) of 100 ms (Nakano et al., 2013). The value of μ_peak_ is different for every ion channel and dictates the maximum modulation attainable during modulation. The modulation factors code for a fractional change in ion channel conductance or in some cases for absolute changes in activation parameter values of ion channels over time. Peak values of modulation factors are shown in Table 2 for dMSN and in Table 3 for iMSN.

**Table 2:** Parameters for dopamine modulation factors - dMSN.

| *Channel $z$* | *Peak value of modulation factor ($\mu_{peak, z}$)* |
| --- | --- |
| *NaF* | 0.95 |
| *Kir* | 1.25 |
| *CaL1.2* | 2.0 |
| *CaN* | 0.2 |
| *CaQ* | 0.5 |
| *NMDA* | 1.3 |

**Table 3:** Parameters for dopamine modulation factors - iMSN.

| *Channel $z$* | *Peak value of modulation factor ($\mu_{peak, z}$)* |
| --- | --- |
| *NaF* | 1.1 |
| *KAs* | 1.1 |
| *CaL1.3* | 0.75 |
| *AMPA* | 0.8 |

Apart from these, the ‘m’ activation parameter of CaL1.3 shifts by −10 mV in dMSN and the ‘h’ inactivation parameter of NaF shifts by +3 mV in iMSN at peak response via dopamine modulation factor (Moyer et al., 2007; Nakano et al., 2013). In Fig. 1, a schematic representation of all the ion channels that were modulated in the spines and the dendritic segment are shown.

**Figure 1.**
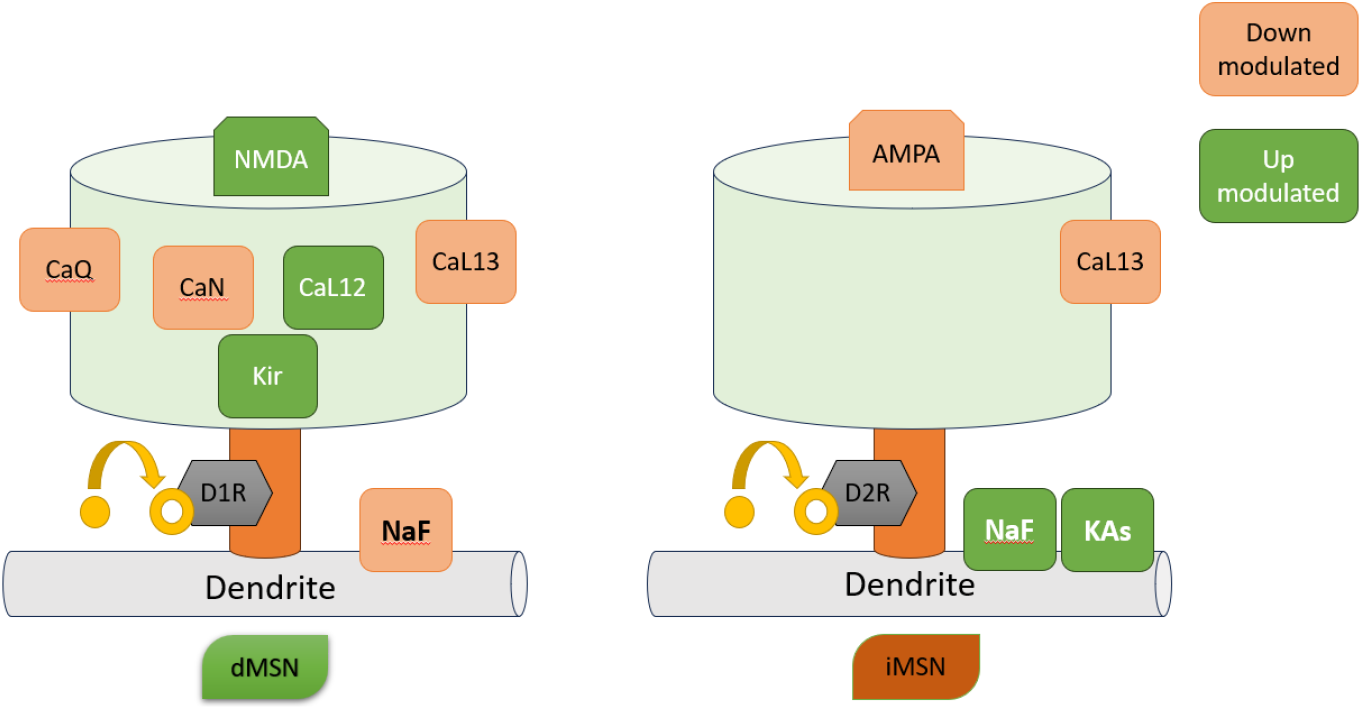
Ion channels modulated by D1R on dMSN (left) and by D2R on iMSN (right). In both MSN sub-types, if the channels are shown in green, then they are up-modulated and if the channels are shown in orange, then they are down-modulated

#### Local dendritic dopamine mechanism in our models

Although dopamine receptors can be present on soma and dendrites, they are predominantly expressed on the neck region of dendritic spines (Yao et al., 2008). Furthermore, NaF channels are not present on dendritic spines but they are reported to be modulated, differentially, by both D1R and D2R. This modulation may arise from the action of dopamine receptors on the NaF channels present on dendritic shaft. Due to volume transmission, it is likely that the DA receptors located on the dendritic shaft are activated along with those located on the dendritic spines, thereby explaining the modulation of NaF channels. From these considerations, we have added DA receptors to the neck section of all dendritic spines and on the dendritic shafts where spines are present, i.e the middle and distal dendrites. Thus, when dopamine input arrives at a dendritic spine, DA receptors on both spine and the localized segment of dendritic shaft are activated (Fig. 2).

**Figure 2.**
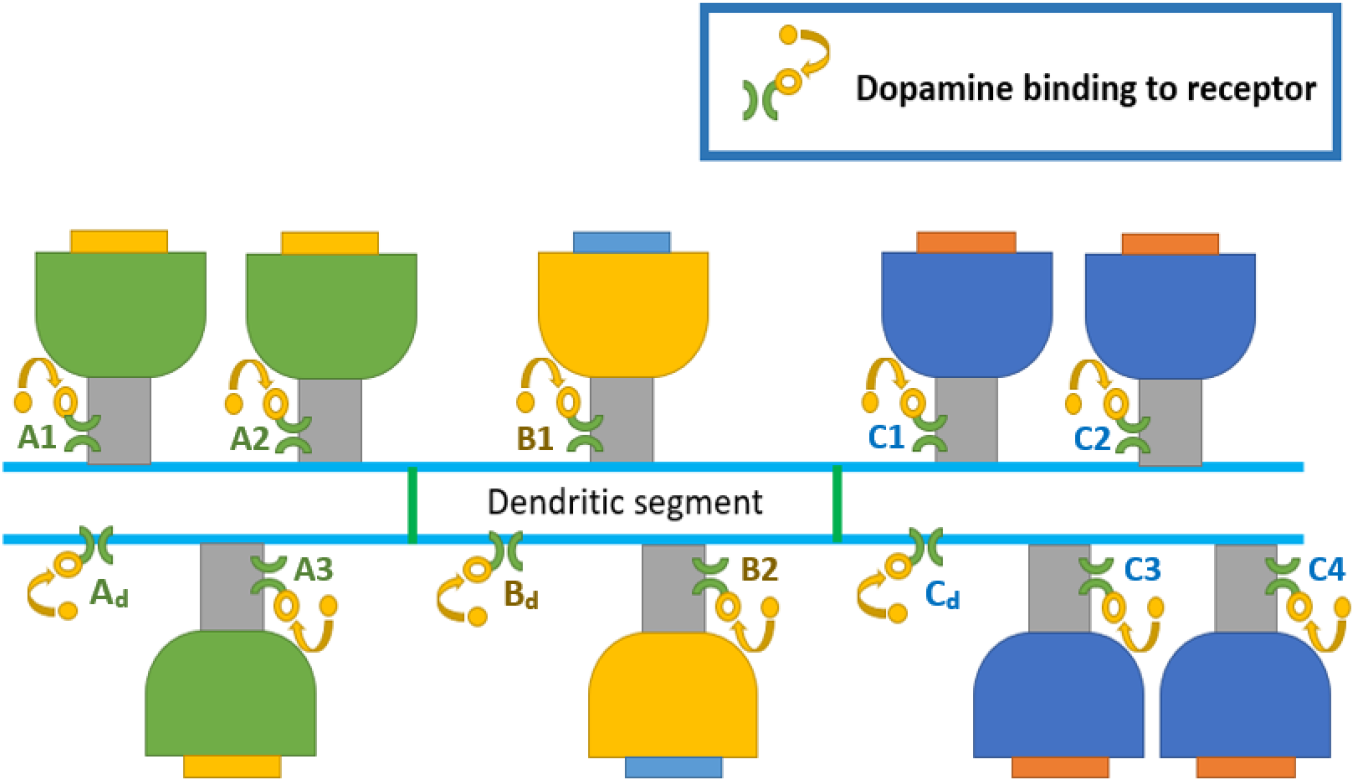
Dopamine modulation of spines and the local dendritic segment. When dopamine receptors labelled A1, A2 or A3 are activated then dendritic dopamine receptor labelled as A_d_ is concurrently activated. Similarly, B_d_ is concurrently activated with B1 or B2, and Cd is concurrently activated with C1, C2, C3 or C4 activation.

#### Method for calculating the % receptor activation

To quantitatively estimate the % activation of dopamine receptors at a given point in time in the model, we first considered the duration that the modulation factor is active (i.e., 60 ms) during which there is a rise in DA modulation activity due to DA binding (Nakano et al., 2013), (Dreyer et al., 2010). If N_act_ is the number of activated dopamine receptor locations and ‘f’ is frequency of dopamine input, then N_act_*f/1000 is number of activations per ms over all active locations. Considering this along with the duration of modulation factor activation we get, (N_act_/N_tot_)*(f/1000)*60 as the fraction of activations where N_tot_ is the total number of dopamine receptors present in the model. Therefore, % activation (at any given instant) = 6*f* (N_act_/N_tot_)

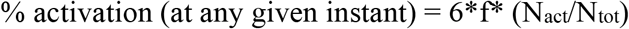

Since DA undergoes volume transmission, the spread of DA can be spatially extensive, making it difficult to decide which receptors to exclude from activation. Accordingly, all dendritic spines were considered for activation by DA although the timing of activation of these receptors at each spine location was different to account for variability in the arrival DA at various locations. A few preliminary studies were also conducted with varying number of spine locations activated; it was inferred from these that the effect of DA receptor activation was greater only on spines where the glutamatergic inputs were also active. Thus, we have activated all spine locations (N_act_ = N_tot_) in this study and only varied the frequency of dopamine inputs, making % activation = 6*f at any given instant. Consequently, the % activation values were taken as multiples of 6.

In this study, the five % receptor activation conditions were: 6%, 24%, 36%, 48% and 72%. Although the middle 3 conditions are evenly spaced, on either ends we have more extreme conditions representing abnormally low or elevated DA receptor activations.

### Interpreting state transition times and dwell time

When there is an increase in glutamatergic inputs (particularly due to arrival of hippocampal inputs) to the cell from 3 Hz to 7.5 Hz or vice-versa, the cell switches its stability by transitioning from a down-state to an up-state or vice-versa (Wolf et al., 2005). These transitions between states are not instantaneous and is gradually driven by changes in activity of several ion channels. Furthermore, GABAergic inputs represent inputs from interneurons and lateral connections from other MSN and the ratio of GABAergic to glutamatergic inputs was maintained constant (∼1:1) during the simulation for both cell-types (Blackwell et al., 2003).

To calculate the transition time, we measure the time taken from ‘input frequency switch’ till it reaches a threshold voltage (not synonymous to AP voltage threshold as explained later).

Henceforth, we refer the down-state to up-state transition time as ‘*upward transition time’* and the up-state to down-state transition time as ‘*downward transition time’*. This threshold to indicate a successful transition was considered to be 5% below the mean up-state voltage for upward transition and 5% above the mean down-state voltage for downward transition. The mean state voltages for dMSN and iMSN were calculated previously over several simulations and found to be significantly different, therefore different threshold values were considered for both cell types as shown in Table 4.

**Table 4:** Threshold voltage (in mV) for calculating state transition times.

| **Cell-type** | **up-state to down-state** | **down-state to up-state** |
| --- | --- | --- |
| dMSN | -70 | -58 |
| iMSN | -68.5 | -60 |

Next, we formulated a parameter called *‘up-state dwell time’* that calculates the actual up-state duration by accounting for transition times using a standardized formula. This is different from ‘state duration’ that is typically considered as 284 ms in literature i.e. duration of glutamatergic input frequency (Blackwell et al., 2003). With consideration to a typical simulation profile, as shown in figure 3, the up-state dwell time (t_dw_) can be calculated as:

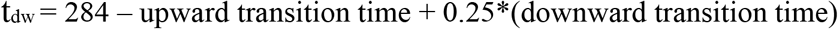

**Figure 3.**
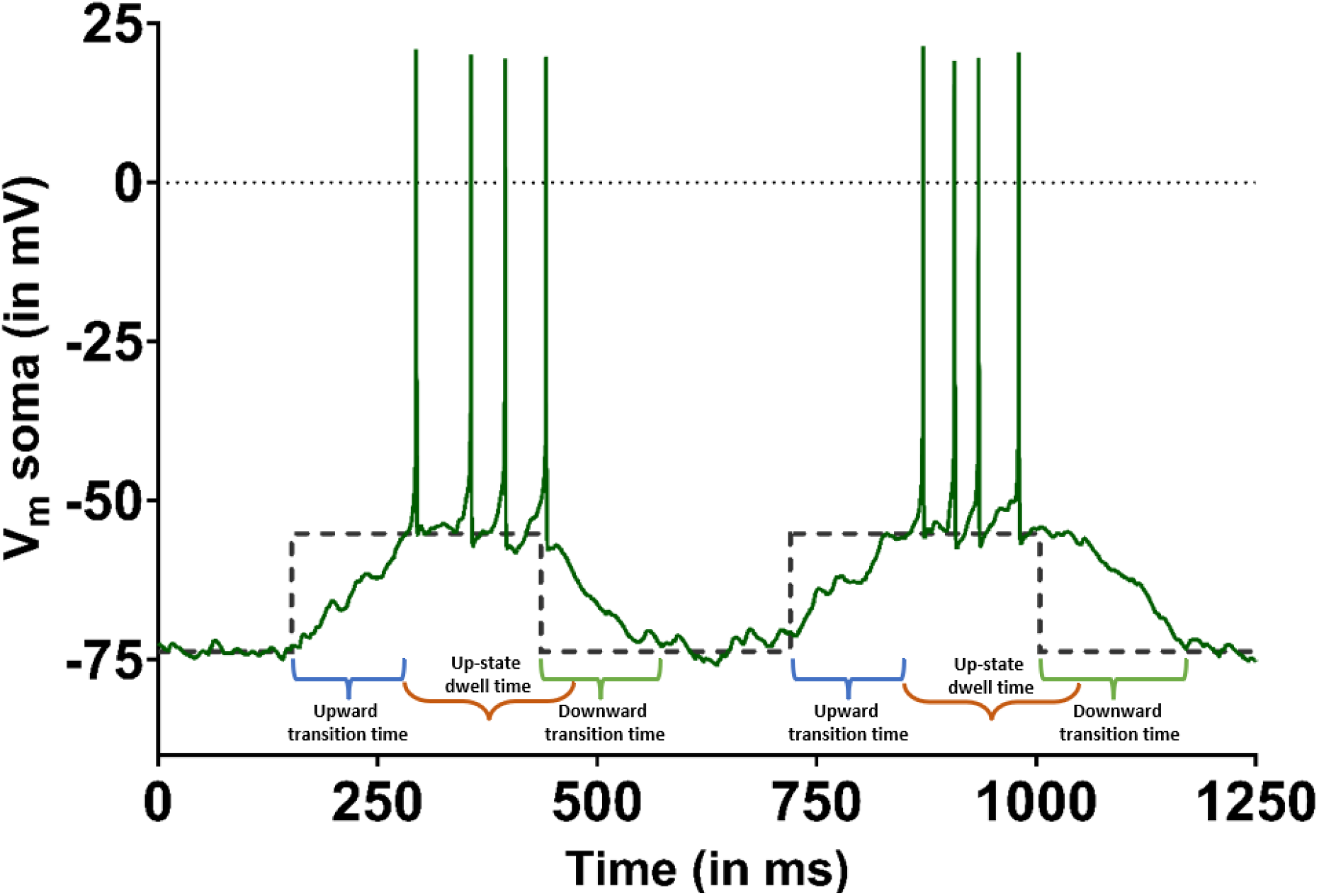
Sample simulation of dMSN model showing upward (down-state to up-state) transition, up-state dwell time and downward (up-state to down-state) transition (dashed black lines indicate switching of glutamatergic input frequency depicted by corresponding mean down-state voltage and mean up-state voltage)

## Results

### Validation of dMSN and iMSN models: Current injection protocol

As reference points to validate our models, experimental data are available for excitability profiles under whole cell current clamp recordings for MSN from the core region of rat Nucleus accumbens (Wolf et al., 2005). Since this is the specific region and species of interest, we validated our models against these. The data do not indicate whether they are for dMSN or iMSN but we can infer that they correspond to dMSN as the morphology and biophysics closely match this sub-type. For example, the total dendritic length reported from experiments for dMSN (3283 μm), matches closely with the (Wolf et al., 2005) model’s total dendritic length of 3280 μm.

It is well established that the intrinsic excitability of dMSN is lower than that of iMSN irrespective of the organism or region (Gertler et al., 2008; Planert et al., 2013; Al-Muhtasib et al., 2018). In accordance with this observation, our iMSN model exhibited greater excitability as expected from the lower total dendritic length and presence of partially inactivating Kir channels (Gertler et al., 2008; Steephen and Manchanda, 2009). The models were injected with step currents of duration 500 ms and varying strengths, matching the experimental protocol, to estimate the intrinsic excitability and compare it with experimental data.

Models replicated the response to current injection as reported in experiments (Fig. 4). In response to negative current injection, the extents of hyperpolarization observed in dMSN (−102.2 mV) and iMSN (−109.2 mV) were different (Fig 4(a) and 4(b)). We also explored strength-duration curves, as an index of excitability, used for the generation of a single spike at different durations. Spikes in iMSN were generated at lower amplitudes of injected current at all durations, indicating their higher excitability compared with dMSN. Moreover, dMSN exhibited a higher rheobase current of 244 pA compared to iMSN with rheobase current of 174 pA (Fig 4(c)). Next, in order to determine the input resistance (R_in_) from the current-voltage graph we observe the sub-threshold voltage responses generated after injecting step currents for 500 ms. We have considered the linear region of sub-threshold responses to determine the R_in_ for the cells. The values of R_in_ were determined to be 82.25 MΩ for dMSN and 120.88 MΩ, corroborating the lower excitability for dMSN than for iMSN (Fig 4(d)).

**Figure 4:**
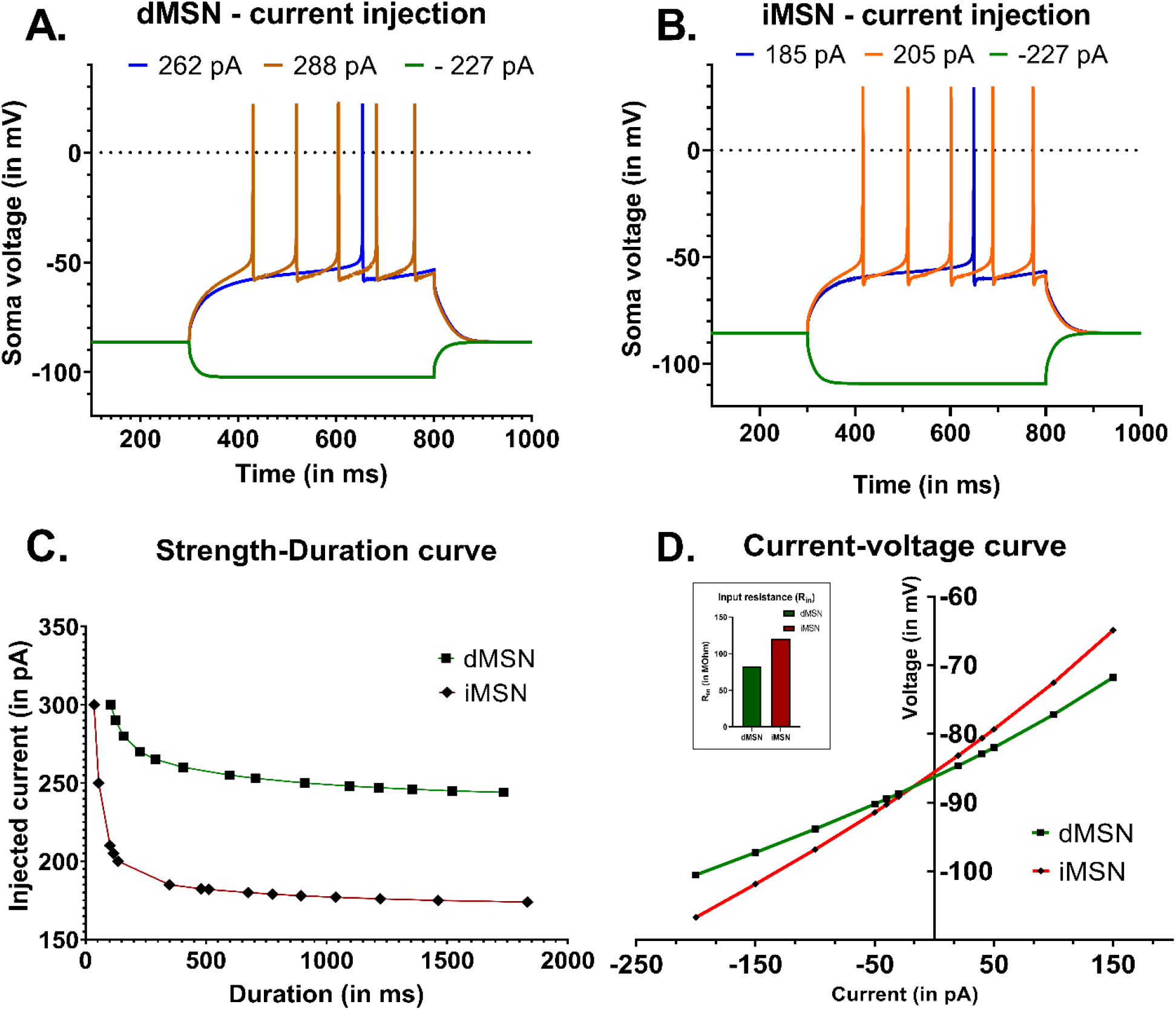
(A) Responses of dMSN to injected step currents (values indicated in legend) of 500 ms duration (B) Responses of iMSN to injected step currents (values indicated in legend) of 500 ms duration. (C) Strength-duration curves for dMSN and iMSN indicate the greater excitability of iMSN over dMSN as less current is needed to generate a spike at each duration of time. (D) Current-voltage curves for dMSN and iMSN models. They show the saturation of voltage at different sub-threshold currents injected for 500 ms in soma. Voltage was measured at the peak of saturation i.e 500 ms. Inset, Input resistance (Rin) of the MSN sub-types were determined to be 82.25 MΩ for dMSN and 120.875 MΩ for iMSN.

### Effect of dopamine on state transition times and up-state dwell time in dMSN and iMSN

The upward transition time and downward transition time (as described in Methods) were calculated for dMSN and iMSN under increasing % activation of D1R and D2R, respectively. For each simulation, there were 6 upward transitions and 6 downward transitions; and for each % receptor activation condition, there were 10 simulations, thereby giving 60 data points for each condition. Consecutive activation conditions were compared using a paired t-test and to correct the familywise error rates from multiple comparisons, Holm-Bonferroni method was used. The p-values and corrected α values were tabulated for upward transition time in Table 5 and for downward transition time in Table 6. In Fig. 5(a) and 5(b), it can be observed that for dMSN, the upward transitions became more rapid while the downward transitions became slower. However, for iMSN we observe the opposite where the upward state transitions got slower and the downward state transitions got faster.

**Table 5:** Paired t-test for Upward transition time: (Down-state to Up-state)

|  | %D1R activation for<br>*dMSN* |  | %D2R activation for<br>*iMSN* |  |
| --- | --- | --- | --- | --- |
| **Comparison** | p-values | corrected $\alpha$ (Rank) | p-values | corrected $\alpha$ (Rank) |
| 6% 24% | 0.0008 | 0.025 (#3) | <0.0001 | 0.025 (#3) |
| 24% 36% | 0.0005 | 0.0167 (#2) | <0.0001 | 0.0125 (#1) |
| 36% 48% | 0.0368 | 0.05 (#4) | 0.0008 | 0.05 (#4) |
| 48% 72% | <0.0001 | 0.0125 (#1) | <0.0001 | 0.0167 (#2) |

**Table 6:** Paired t-test for Downward transition time: (Up-state to Down-state)

|  | %D1R activation for<br>*dMSN* |  | %D2R activation for<br>*iMSN* |  |
| --- | --- | --- | --- | --- |
| **Comparison** | p-values | corrected $\alpha$ (Rank) | p-values | corrected $\alpha$ (Rank) |
| 6% 24% | <0.0001 | 0.0167 (#2) | 0.0096 | 0.0167 (#2) |
| 24% 36% | <0.0001 | 0.0125 (#1) | 0.0116 | 0.025 (#3) |
| 36% 48% | 0.1654 | 0.05 (#4) | 0.2702 | 0.05 (#4) |
| 48% 72% | 0.0001 | 0.025 (#3) | 0.0091 | 0.0125 (#1) |

**Figure 5.**
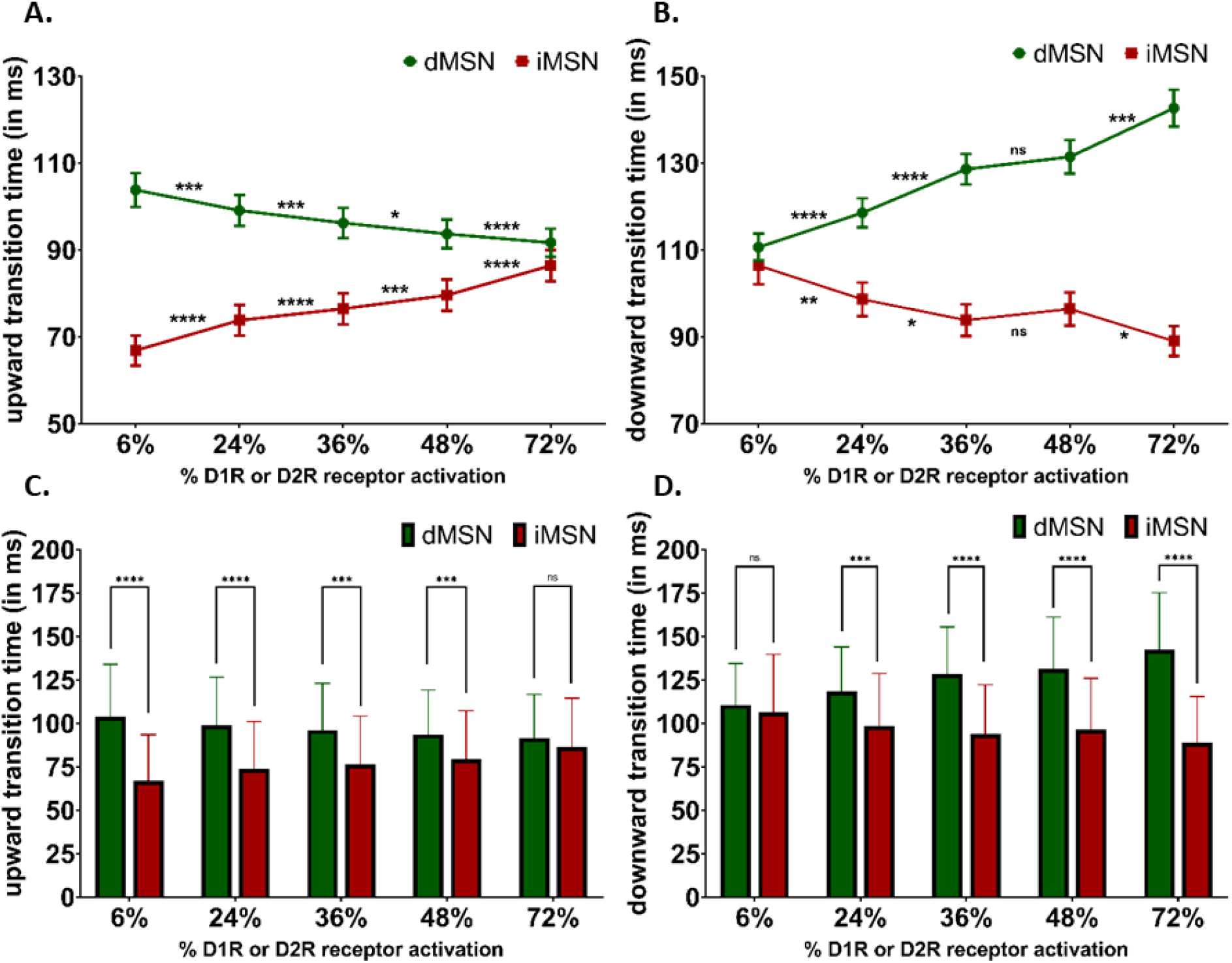
A) Upward transition times and B) Downward transition times compared across increasing % DA receptor activation C) Upward transition times and D) Downward transition times compared between dMSN (left) and iMSN (right) for different % DA receptor activation (n = 60)

**Figure 6.**
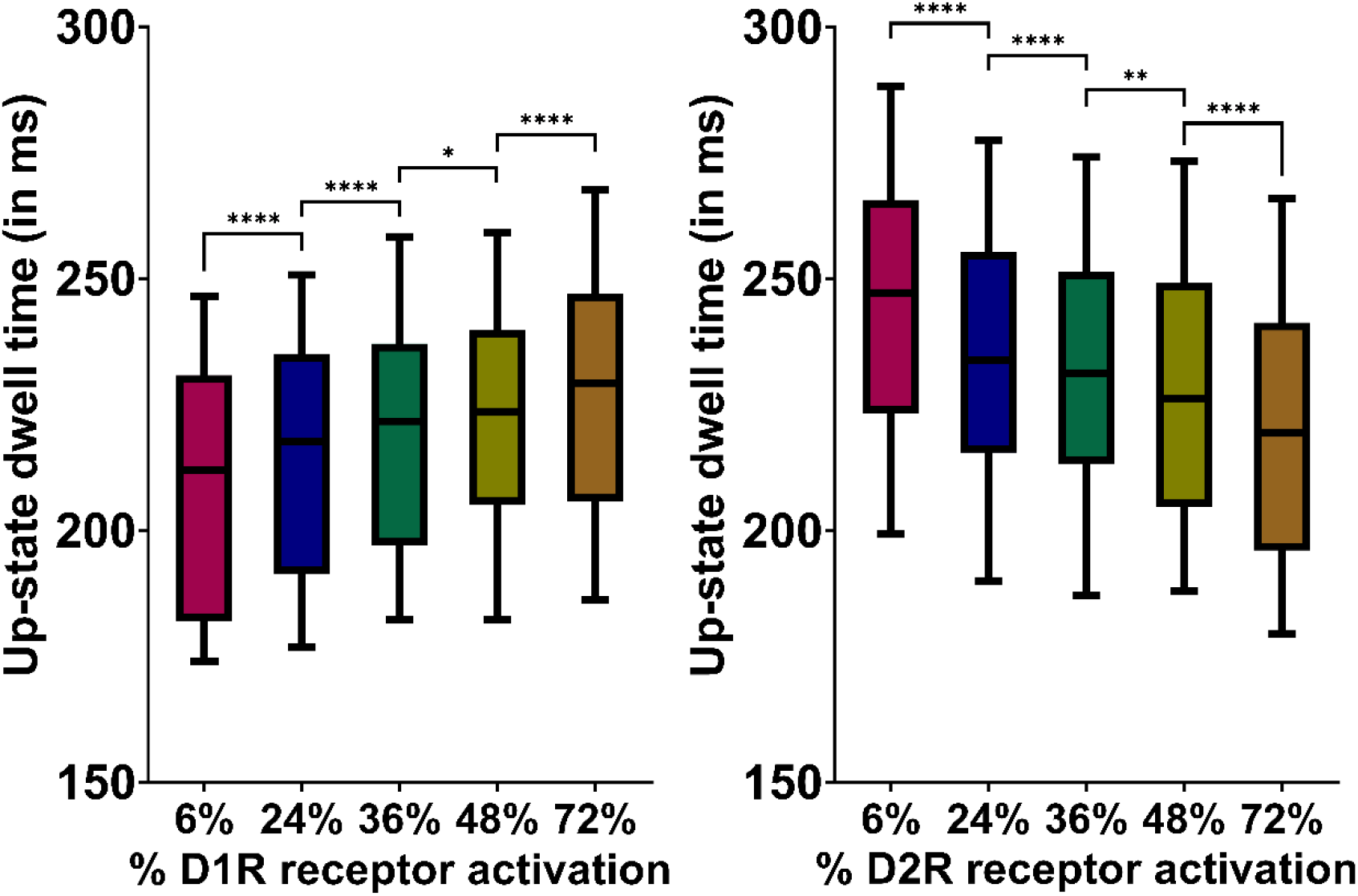
Up-state dwell time across increasing % DA receptor activation conditions increased for dMSN (left) and decreased for iMSN (right).

**Figure 7.**
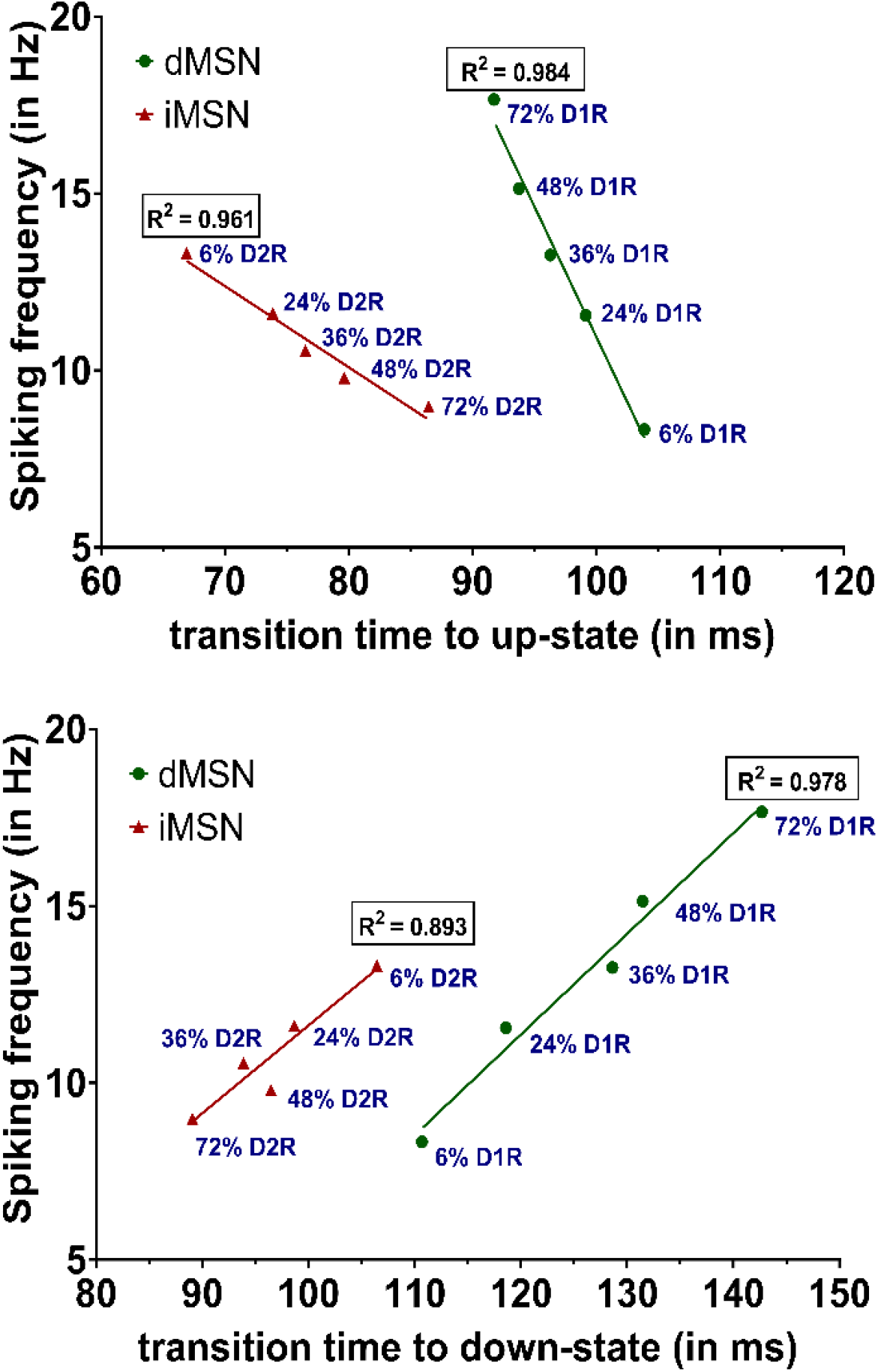
Correlation between the mean spiking frequency and mean upward transition time (top) and downward transition time (bottom) for dMSN (green) and iMSN (red)

In Fig. 5(c) and 5(d), State transition times of dMSN and iMSN were compared at each % DA receptor activation. The upward transition times for dMSN and iMSN were significantly different at lower % DA receptor activation but they got more similar as the % activation was increased. On the other hand, the downward transition time was similar at low % receptor activation but became increasingly different with increasing activation.

Next, the up-state dwell time was analysed. This depends on both the upward transition and downward transition as explained in Methods section. With this unified measure, we analyse how it changes with increasing % DA receptor activation using paired t-test followed by Holm-Bonferroni method for post hoc correction of α (Table 7). As anticipated, with increasing % receptor activation, dMSN dwell time is greater in the up-state whereas iMSN dwell time is lower in the up-state. However, in absolute terms, the highest mean residence time for dMSN at 72% activation was still lower than that of iMSN across all conditions except 72% activation.

**Table 7:**
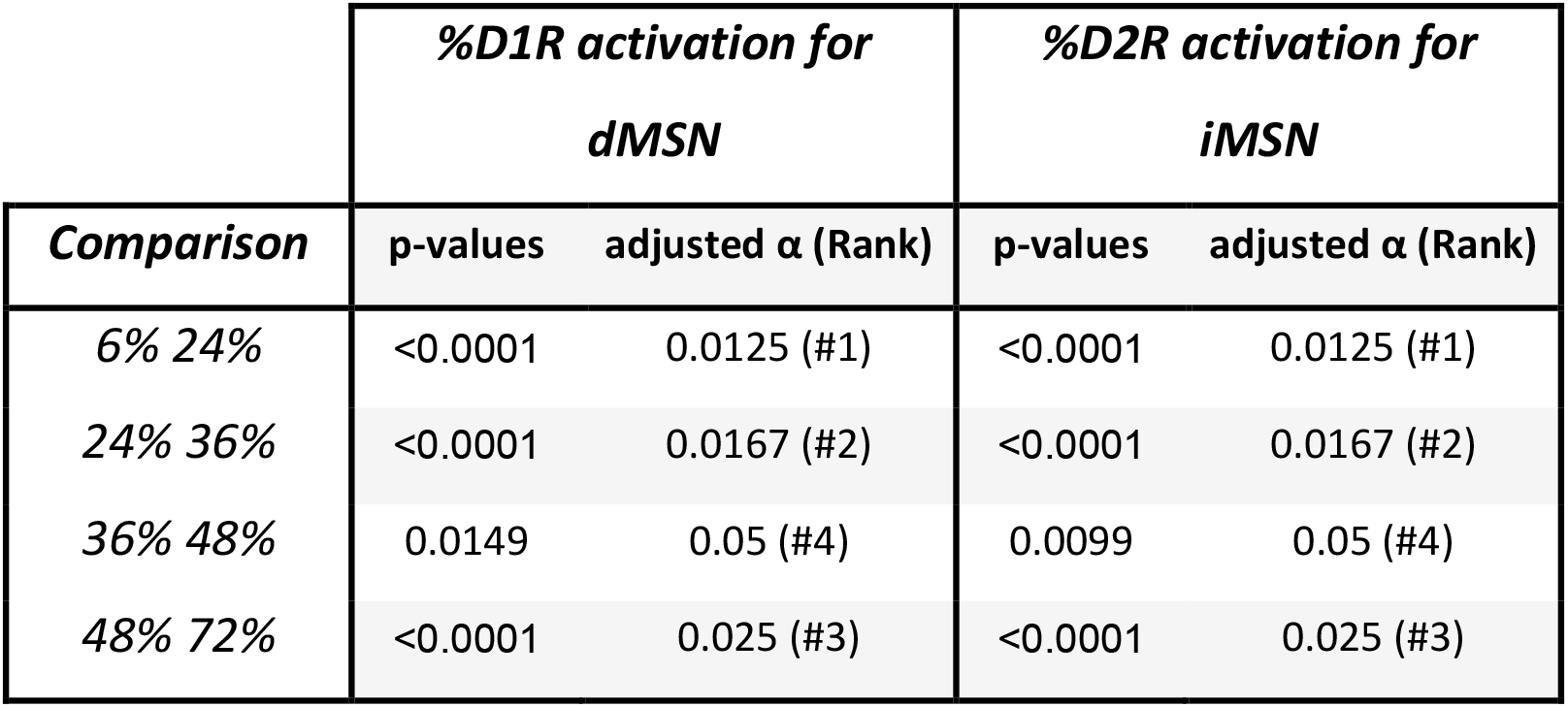
Hypothesis comparison for dwell time across % receptor activation.

### Correlation of state transition times with spiking frequency

For our range of % activation conditions, we analyse the correlation between the mean spiking frequency and mean transition times for dMSN and iMSN. A strong correlation (R^2^ > 0.85) was observed between transition times and spiking frequency for all % activation conditions. The effect of increasing % DA receptor activation was opposite for dMSN and iMSN, as expected. Additionally, it was observed that the changes in spiking frequency occupied a wider range of values for dMSN than iMSN, indicating greater sensitivity to DA receptor activation. Although there is strong correlation, lower transition time does not always translate to higher spiking frequency. For example, despite having slower upward transition times, dMSN at 48% or higher D1R activation had greater spiking frequency than iMSN even under its most excitable conditions.

## Discussion

Like the Ship of Theseus analogy, although the biophysically constrained models used for this study can be traced back to (Wolf et al., 2005), it is difficult to say whether the models are derived considering how several modifications were incorporated. The code was written in Python and NEURON simulation environment was used to run the simulations. The methods section focused on newer details such as the dendritic spines and explicit spatiotemporal modulation of ion channels through dopamine (DA) receptors. A crucial addition is the localized dopaminergic ion channel modulation on the dendrites. The mechanism works such that when DA receptor activates on a dendritic spine, the DA receptor on the corresponding small segment of distal dendrite also activates. This explains how ion channels, such as NaF or KAs, that are absent in the dendritic spine but present on the dendritic segment are modulated.

For validating the models, current injection data were not available for the excitability profile of iMSN of rat nucleus accumbens core region. However, evidence from patch-clamp recording from dorsal striatum of rats indicates that the rheobase current (mean ± SD) for dMSN was 191.42 ± 67.36 pA and for iMSN was 139.1 ± 49.58 pA (Planert et al., 2013). Furthermore, whole-cell current-clamp recordings of dMSN in rat nucleus accumbens core region indicates rheobase currents in the range of 200 pA to 300 pA (Wolf et al., 2005). Collating these data, it is reasonable to expect that the excitability profiles of MSN sub-types from dorsal or ventral striatum of rat will be similar, although with rheobase current of iMSN being lower than dMSN due to greater excitability of the former. Therefore, the values obtained (257.6 pA for dMSN and 182.5 pA for iMSN) fall within acceptable range of MSN excitability. Additionally, dMSN contain the neuropeptide dynorphin and substance P (SP), thereby they would express non-inactivating Kir channels whereas iMSN contains inactivating Kir channels (Steephen and Manchanda, 2009; Humphries and Prescott, 2010). This could be one reason for varied response of dMSN and iMSN for hyperpolarizing current injection.

The transition times were compared across consecutive % DA receptor activation. This led to multiple comparisons even though we are not comparing all possible pairs. Consequently, based on preliminary exploration, paired t-tests were followed by post-hoc Holm-Bonferroni correction in all necessary cases. With increasing activation of dopamine receptors, dMSN transitions to the up-state were accelerated while their transition to the down-states were unhurried. This suggests the influence of increasing DA receptor activation to keep dMSN in up-state for longer duration. Furthermore, for iMSN, the opposite was observed wherein the transitions to up-state got slower while the transitions to downstate were speeded up. This was corroborated by our findings on the up-state dwell time. For dMSN, it was increasing whereas for iMSN it was decreasing. An interesting observation is that DA is able to alter state transition times even with the glutamatergic input frequencies, which are key to occurrence of up-states and down-states, remaining constant.

In absolute terms, dMSN mean state transition times were slower than that of iMSN for all conditions and irrespective of the direction of transition. This may be attributed to the fact that iMSN expresses partially inactivating Kir channels whose inactivation facilitates departure from the down-state and shortening the absolute time of the transitions. On the other hand, the dMSN expresses non-inactivating Kir channels that tend to hold the cell in the hyperpolarized down-state. Theoretically, as spikes occur only during the up-state, the longer the cell dwells in the up-state, the greater is the window for more spiking to occur. However, there could be other factors wherein even if the up-state dwell time increases, the inter-spike interval can also increase, leading to little or no change in spiking frequency.

Considering the integral part that MSN sub-types play in driving the direct and indirect pathways in the basal ganglia, detailed biophysical models as described here can be quite useful in uncovering the functioning of the reward circuitry. While this research laid foundation to the biophysical models for dMSN and iMSN, in our follow-up research paper, we tried to uncover how this transient dopamine, acting as a motivational signal, interacts with MSN sub-types under different bias conditions to influence action selection framework towards exploration over exploitation of actions.

## References

Al-Muhtasib, N., Forcelli, P.A., Vicini, S., 2018. Differential electrophysiological properties of D1 and D2 spiny projection neurons in the mouse nucleus accumbens core. Physiological Reports. 6, e13784.

Blackwell, K.T., Czubayko, U., Plenz, D., 2003. Quantitative estimate of synaptic inputs to striatal neurons during up and down states in vitro. J Neurosci. 23, 9123–32.

Dreyer, J.K., et al., 2010. Influence of phasic and tonic dopamine release on receptor activation. J Neurosci. 30, 14273–83.

Gangarossa, G., et al., 2013. Distribution and compartmental organization of GABAergic medium-sized spiny neurons in the mouse nucleus accumbens. Front Neural Circuits. 7, 22.

Gerfen, C.R., Surmeier, D.J., 2011. Modulation of striatal projection systems by dopamine. Annu Rev Neurosci. 34, 441–66.

Gertler, T.S., Chan, C.S., Surmeier, D.J., 2008. Dichotomous anatomical properties of adult striatal medium spiny neurons. J Neurosci. 28, 10814–24.

Grace, A.A., 2000. Gating of information flow within the limbic system and the pathophysiology of schizophrenia. Brain Res Brain Res Rev. 31, 330–41.

Gruber, A.J., et al., 2003. Modulation of striatal single units by expected reward: a spiny neuron model displaying dopamine-induced bistability. J Neurophysiol. 90, 1095–114.

Gurney, K.N., Humphries, M.D., Redgrave, P., 2015. A new framework for cortico-striatal plasticity: behavioural theory meets in vitro data at the reinforcement-action interface. PLoS Biol. 13, e1002034.

Hansen, K.B., Yuan, H., Traynelis, S.F., 2007. Structural aspects of AMPA receptor activation, desensitization and deactivation. Curr Opin Neurobiol. 17, 281–8.

Humphries, M.D., Prescott, T.J., 2010. The ventral basal ganglia, a selection mechanism at the crossroads of space, strategy, and reward. Prog Neurobiol. 90, 385–417.

Kolibius, L.D., et al., 2023. Hippocampal neurons code individual episodic memories in humans. Nat Hum Behav. 7, 1968–1979.

Lau, B., Monteiro, T., Paton, J.J., 2017. The many worlds hypothesis of dopamine prediction error: implications of a parallel circuit architecture in the basal ganglia. Curr Opin Neurobiol. 46, 241–247.

Le Moine, C., Bloch, B., 1995. D1 and D2 dopamine receptor gene expression in the rat striatum: sensitive cRNA probes demonstrate prominent segregation of D1 and D2 mRNAs in distinct neuronal populations of the dorsal and ventral striatum. J Comp Neurol. 355, 418–26.

Li, Y.X., Rinzel, J., 1994. Equations for InsP3 receptor-mediated [Ca2+]i oscillations derived from a detailed kinetic model: a Hodgkin-Huxley like formalism. J Theor Biol. 166, 461–73.

Li, Z., et al., 2018. Cell-Type-Specific Afferent Innervation of the Nucleus Accumbens Core and Shell. Front Neuroanat. 12, 84.

Lindroos, R., et al., 2018. Basal Ganglia Neuromodulation Over Multiple Temporal and Structural Scales-Simulations of Direct Pathway MSNs Investigate the Fast Onset of Dopaminergic Effects and Predict the Role of Kv4.2. Front Neural Circuits. 12, 3.

Mattioni, M., Le Novere, N., 2013. Integration of biochemical and electrical signaling-multiscale model of the medium spiny neuron of the striatum. PLoS One. 8, e66811.

Moyer, J.T., Wolf, J.A., Finkel, L.H., 2007. Effects of dopaminergic modulation on the integrative properties of the ventral striatal medium spiny neuron. J Neurophysiol. 98, 3731–48.

Nakano, T., Yoshimoto, J., Doya, K., 2013. A model-based prediction of the calcium responses in the striatal synaptic spines depending on the timing of cortical and dopaminergic inputs and post-synaptic spikes. Frontiers Computational Neuroscience. 7, 119.

Nicola, S.M., Surmeier, J., Malenka, R.C., 2000. Dopaminergic modulation of neuronal excitability in the striatum and nucleus accumbens. Annu Rev Neurosci. 23, 185–215.

O’Donnell, P., Grace, A.A., 1995. Synaptic interactions among excitatory afferents to nucleus accumbens neurons: hippocampal gating of prefrontal cortical input. J Neurosci. 15, 3622–39.

Planert, H., Berger, T.K., Silberberg, G., 2013. Membrane properties of striatal direct and indirect pathway neurons in mouse and rat slices and their modulation by dopamine. PLoS One. 8, e57054.

Rane, M., Manchanda, R., 2017. Effect of Spine Density on Excitability in Accumbal Medium Spiny Neurons-A Computational Approach. Journal of Addiction Research & Therapy. 08.

Steephen, J.E., Manchanda, R., 2009. Differences in biophysical properties of nucleus accumbens medium spiny neurons emerging from inactivation of inward rectifying potassium currents. J Comput Neurosci. 27, 453–70.

Wilson, C.J., et al., 1983. Three-dimensional structure of dendritic spines in the rat neostriatum. J Neurosci. 3, 383–8.

Wilson, C.J., Kawaguchi, Y., 1996. The origins of two-state spontaneous membrane potential fluctuations of neostriatal spiny neurons. J Neurosci. 16, 2397–410.

Wolf, J.A., et al., 2005. NMDA/AMPA ratio impacts state transitions and entrainment to oscillations in a computational model of the nucleus accumbens medium spiny projection neuron. J Neurosci. 25, 9080–95.

Yao, W.D., Spealman, R.D., Zhang, J., 2008. Dopaminergic signaling in dendritic spines. Biochem Pharmacol. 75, 2055–69.

